# Evaluating vaccine-elicited antibody activities against *Neisseria gonorrhoeae:* cross-protective responses elicited by the 4CMenB meningococcal vaccine

**DOI:** 10.1101/2023.08.03.551882

**Authors:** Mary C. Gray, Keena S. Thomas, Evan R. Lamb, Lacie M. Werner, Kristie L. Connolly, Ann E. Jerse, Alison K. Criss

## Abstract

The bacterial pathogen *Neisseria gonorrhoeae* is an urgent global health problem due to increasing numbers of infections, coupled with rampant antibiotic resistance. Vaccines against gonorrhea are being prioritized to combat drug-resistant *N. gonorrhoeae.* Meningococcal serogroup B vaccines such as 4CMenB are predicted by epidemiology studies to cross-protect individuals from natural infection with *N. gonorrhoeae* and elicit antibodies that cross-react with *N. gonorrhoeae.* Evaluation of vaccine candidates for gonorrhea requires a suite of assays for predicting efficacy in vitro and in animal models of infection, including the role of antibodies elicited by immunization. Here we present assays to evaluate antibody functionality after immunization: antibody binding to intact *N. gonorrhoeae,* serum bactericidal activity, and opsonophagocytic killing activity using primary human neutrophils (polymorphonuclear leukocytes). These assays were developed with purified antibodies against *N. gonorrhoeae* and used to evaluate serum from mice that were vaccinated with 4CMenB or given alum as a negative control. Results from these assays will help prioritize gonorrhea vaccine candidates for advanced preclinical to early clinical study and will contribute to identifying correlates and mechanisms of immune protection against *N. gonorrhoeae*.

## Introduction

Gonorrhea is a prominent global health threat, with 82.4 million cases estimated worldwide in 2020 (1). Challenges to the control of gonorrhea include increasing numbers of cases in the United States and worldwide, the high percentage of asymptomatic infections, and limited treatment options due to increasing resistance to the antibiotics used to treat gonorrhea (1–3). Untreated gonorrhea or where there is treatment failure has significant consequences for human health, including, pelvic inflammatory disease leading to chronic pelvic pain, infertility, the life-threatening risk of ectopic pregnancy, and arthritis (4). Moreover, individuals with gonorrhea have an increased risk of both acquiring and spreading other sexually transmitted infections including HIV and chlamydia (5, 6). These issues have prompted organizations including the World Health Organization and US Centers for Disease Control and Prevention to call for the development of vaccines for gonorrhea (3, 7). Epidemiological modeling predicts that even a partially protective vaccine would help reduce the prevalence of gonorrhea, especially in populations experiencing high levels of infection (8–11).

Despite decades of effort, an effective gonorrhea vaccine has not yet been developed. The bacterium that causes gonorrhea, *Neisseria gonorrhoeae*, undergoes extensive phase and antigenic variation, as well as allelic recombination of immunodominant surface structures such as type IV pili, opacity-associated proteins, lipooligosaccharide, and the PorB porin (12). Further, antibodies against the reduction-modifiable protein on the gonococcal surface masks the presentation of PorB to protect it from immune recognition (13). Immunologically, acute gonorrhea is characterized by an inflammatory response in which neutrophils are recruited to sites of mucosal infection, but do not clear the bacteria (14). The sustained neutrophilic inflammatory response may lead to tissue damage rather than control of infection (15). Neither the mechanisms nor the correlates of immune protection for gonorrhea are defined, since infected individuals do not generate protective immunity against subsequent infections (12).

For the related pathogen *Neisseria meningitidis,* bactericidal antibodies are the correlate of protection for vaccines that target the capsular polysaccharide or other surface-exposed components. A similar assumption has been made for *N. gonorrhoeae,* which like *N. meningitidis* is predominantly an extracellular pathogen. At mucosal surfaces, *N. gonorrhoeae* would be exposed to antibodies in secretions and inflammatory serum transudates, and bacteria that translocate to blood for disseminated infection would encounter antibodies in systemic circulation. Antibodies can protect against infection by extracellular pathogens by facilitating complement-mediated deposition on the pathogen surface that leads to lysis and/or opsonophagocytic killing, and also by blocking essential functions of the pathogen such as adherence or nutrient acquisition (16, 17). The complement system is a cascade of highly regulated proteolytic activities that culminate in deposition of C3b on the microbial surface (18). C3b and its degradation products serve as opsonins that bind complement receptors on phagocytic cells to lead to microbial uptake and intracellular killing. C3b is also part of the C5 convertase that is required for formation of the membrane attack complex, a pore comprised of C5b-C9 that leads to microbial rupture. Degradation products C3a, C4a, and C5a increase vascular permeability, and C5a has chemotactic activity for phagocytes, which together enhance immune cell recruitment and activation when encountering microbes. The importance of the complement system in Neisserial pathogenesis is reflected in the ∼1,000-10,000-fold increased susceptibility of individuals with complement deficiencies to invasive meningococcal and gonococcal disease (19).

The introduction of reverse vaccinology and the successful creation and deployment of vaccines against serogroup B *Neisseria meningitidis* have reinvigorated efforts to develop a gonorrhea vaccine. In 2017, immunization with an outer membrane vesicle (OMV)-based meningococcal serogroup B vaccine was reported to reduce the risk of gonorrhea in individuals in New Zealand with an estimated efficacy of 31%, based on a retrospective epidemiological study, without affecting their susceptibility to chlamydia (20). The vaccinated individuals also had a lower hospitalization rate due to gonorrhea (21). Similarly, a different meningococcal B OMV vaccine markedly reduced gonorrhea rates in Cuba by retrospective analysis (22, 23). In 2015, the 4-component meningococcal B vaccine (4CMenB; trade name Bexsero®) was licensed for use in the US in 2015. It is comprised of OMV from the New Zealand serogroup B vaccine strain, along with 3 recombinant antigens (Neisserial adhesin A (NadA), the factor H binding protein fused to the GNA2091 antigen, and Neisserial heparin binding antigen fused to the GNA1030 antigen) (24). These three antigens were included in part due to their ability to elicit antibodies that conferred serum bactericidal activity against serogroup B *N. meningitidis* (25). 4CMenB vaccination reduces the duration of colonization by *N. gonorrhoeae* in a female mouse genital tract infection model (26). Antibodies raised in response to 4CMenB and other meningococcal OMV-based immunogens in mice, rabbits, and humans have been shown to cross-react against antigens in *N. gonorrhoeae* lysates (26–28). Based on these findings, studies have been initiated to determine if 4CMenB confers protection against *N. gonorrhoeae* and gonorrhea in human cohorts. Reports since the New Zealand and Cuba studies suggest a 32-46% reduced incidence of gonorrhea in individuals vaccinated with 4CMenB compared to those who were not, while the incidence of chlamydia in these cohorts was unaffected by vaccination (29–32). Currently, 4CMenB is in several Phase III clinical trials for its efficacy against gonorrhea, and additional OMV-based vaccines, including those with detoxified LPS, are being evaluated preclinically (28). Both defined and novel surface-exposed components of *N. gonorrhoeae,* including protein and lipooligosaccharide antigens, are being evaluated preclinically for their vaccine potential (16, 17, 33–36).

Antibody-mediated serum bactericidal activity (SBA) has long been considered as a correlate of protection for vaccines against *N. meningitidis* (37–40), and SBA titers were used to prioritize antigens for incorporation into 4CMenB (25). While there is less consensus about the contribution of opsonophagocytic killing activity (OPKA) to meningococcal vaccine efficacy, OPKA may help protect individuals whose antibody titer is not sufficient to evoke SBA and in those with terminal complement deficiencies who cannot mount SBA (39, 41–43). Assays to examine SBA and OPKA in *N. gonorrhoeae* are being developed (44, 45). The premise for both SBA and opsonophagocytosis is that vaccine-elicited antibody binds to the surface of bacteria, but this property of vaccination is not typically evaluated in the context of intact pathogens. Instead, isolated antigens are used in enzyme-linked immunosorbent assays (ELISA) or similar in vitro approaches, but the antigens may not be in the conformation, membrane environment, and density as found on the bacterial membrane.

In this report, we present results from three assays to interrogate the functional antibody response against *N. gonorrhoeae* that is elicited by immunization of mice: measurement of antibody binding to the bacterial surface by imaging flow cytometry; SBA assay; and OPKA assay using primary human polymorphonuclear leukocytes (PMNs; predominantly neutrophils). Alternative analyses for complement C3 deposition on *N. gonorrhoeae* and for opsonophagocytosis of *N. gonorrhoeae* by HL-60 promyelocytes using flow cytometry are also presented. As proof of concept, we show that the serum from mice that are vaccinated with meningococcal 4CMenB elicits stronger functional antibody responses than serum from alum-treated mice. Results from these assays will support the quest for a gonorrhea vaccine and will allow the discovery of correlates and mechanisms of immune protection against *N. gonorrhoeae*.

## Materials and Methods

### Bacterial Strains and Growth Conditions

The following *N. gonorrhoeae* strains were used for this study: FA1090 (46), F62 (47), FA19 (48), MS11 (49), and H041 (WHO X) (50, 51). *N. gonorrhoeae* was routinely grown on gonococcal medium base (BD Difco) plus Kellogg’s supplement I and 0.5 mg/ml Fe(NO_3_)_3_ (52) (GCB) plates for 16 hr at 37 °C in 5% CO_2_.

### Human Sera As Complement Sources

For SBA, IgG/IgM-depleted pooled normal human serum (Igd-NHS, PelFreez Catalog #34010, Lot #28341) was used as the complement source. C6-depleted Normal Human Serum (Complement Technology, Catalog #A323) was used as the complement source for OPKA and opsonophagocytosis. Serum was heat-inactivated by incubation at 56 °C for 30 min.

### Antibodies

See **Table 1** for antibodies used in this study.

**Table 1.**
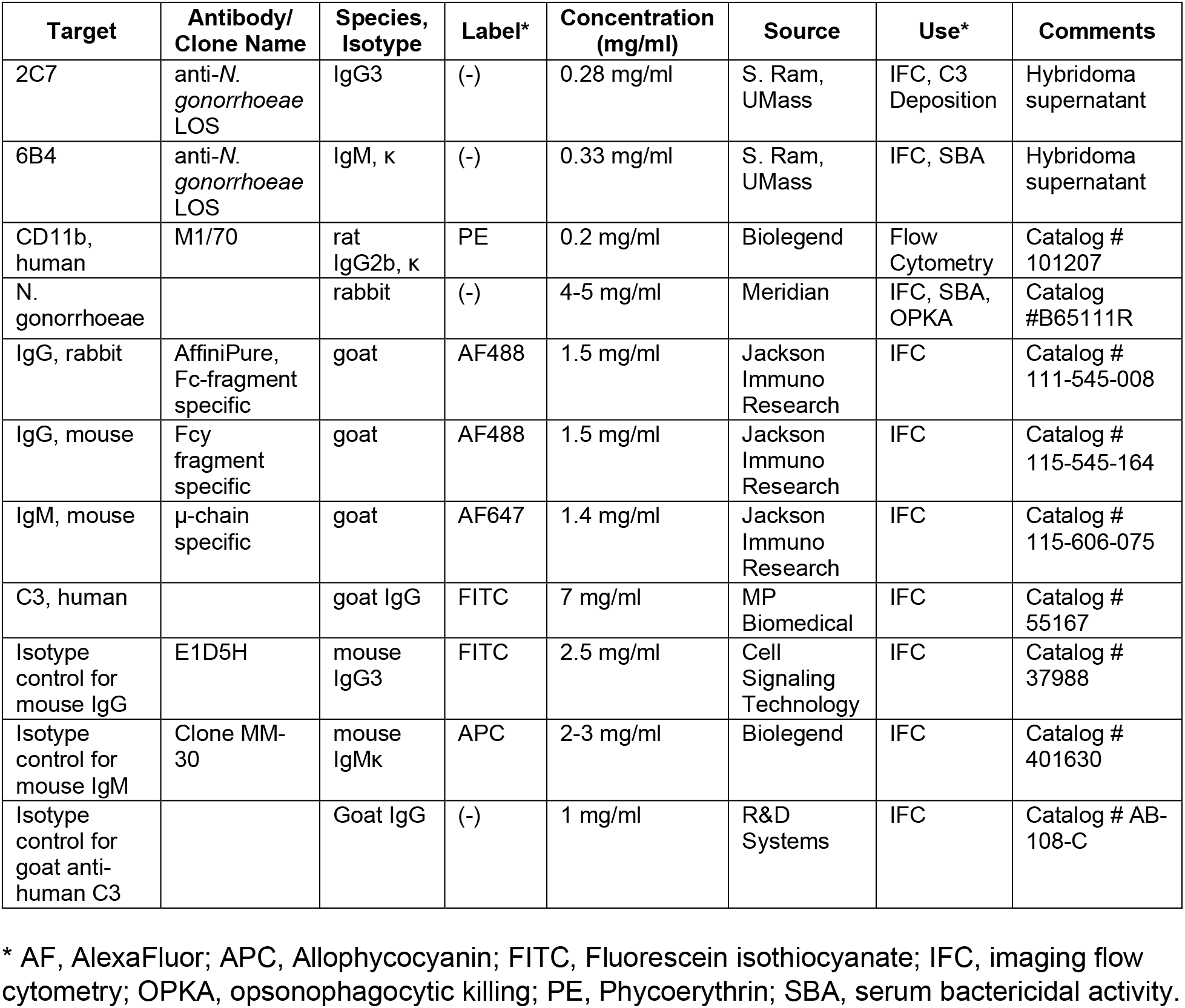
Antibodies used in this study.

### Mouse Sera

All animal experiments were conducted at the Uniformed Services University according to guidelines established by the Association for the Assessment and Accreditation of Laboratory Animal Care using a protocol approved by the University’s Institutional Animal Care and Use Committee. Female Balb/c mice were vaccinated subcutaneously with 250 µl 4CMenB (GlaxoSmithKline) or given an equal volume of alum control. On day 63, sera were collected, pooled, and snap-frozen at −80 °C. Mouse sera were heat-inactivated before use in functional assays. Only female mice were used in preparation for experiments that test the efficacy of vaccine candidates in protecting from experimental genital Gc infection (53).

### Antibody Binding to *N. gonorrhoeae* by Imaging Flow Cytometry

*N. gonorrhoeae* was swabbed from GCB agar into gonococcal base liquid media (GCBL) with Kellogg’s supplement I and 1.25 µM Fe(NO_3_)_3_ • 9 H_2_O (52, 54) to an optical density at 550 nm (OD_550_) of 0.392, pelleted by centrifugation at 10,000 x g for 3 min, and resuspended in 1 ml of RPMI + 2% Bovine Serum Albumin (BSA, heat shock fraction, protease free, essentially globulin free, pH 7, Sigma, Catalog #A3059) to a concentration of ∼2 x 10^8^ CFU/ml. Twenty-five µl of bacterial suspension were added to a 96 well V-bottom plate (Sarstedt, Catalog # 82.1583.001) and incubated with 25 µl of heat-inactivated mouse serum or control monoclonal antibodies (2C7, 6B4) or polyclonal rabbit anti-GC at the indicated concentrations for 30 min at 37 °C, 5% CO_2_. 2C7 was incubated at final concentrations of 70 µg/ml, 28 µg/ml and 14 µg/ml; 6B4 was incubated at final concentrations of 3.3 µg/ml, 1.65 µg/ml and 0.825 µg/ml; and rabbit anti-GC was incubated at final concentrations of 45 µg/ml, 22.5 µg/ml and 11.25 µg/ml. Bacteria were washed 3 times by pelleting in RPMI + 10% Heat-Inactivated Fetal Bovine Serum (HI-FBS). Secondary antibodies were added at a final concentration of 7.5 µg/ml and incubated with bacteria for 30 min at 37 °C, 5% CO_2_. Samples were washed once with PBS, and pellets were resuspended in 2% paraformaldehyde (PFA) containing 5 µg/ml 4’, 6-diamidino-2-phenylindole (DAPI). Samples were analyzed by imaging flow cytometry using Imagestream^X^ Mk II with INSPIRE® software (Luminex Corporation). Alexa Fluor® 488 fluorescence was detected with excitation at 488 nm and emission collected with a 480–560 nm filter. Alexa Fluor® 647 fluorescence was detected with a 658 nm laser excitation and emission was collected with a 660–740 nm filter. DAPI fluorescence was detected with excitation at 405 nm and emission collected with a 420–505 nm filter. Single color samples for Alexa Fluor® 488, Alexa Fluor® 647, and DAPI were collected without brightfield to create a compensation matrix for each experiment. Gating strategy is presented in Figure 1.

**Figure 1.**
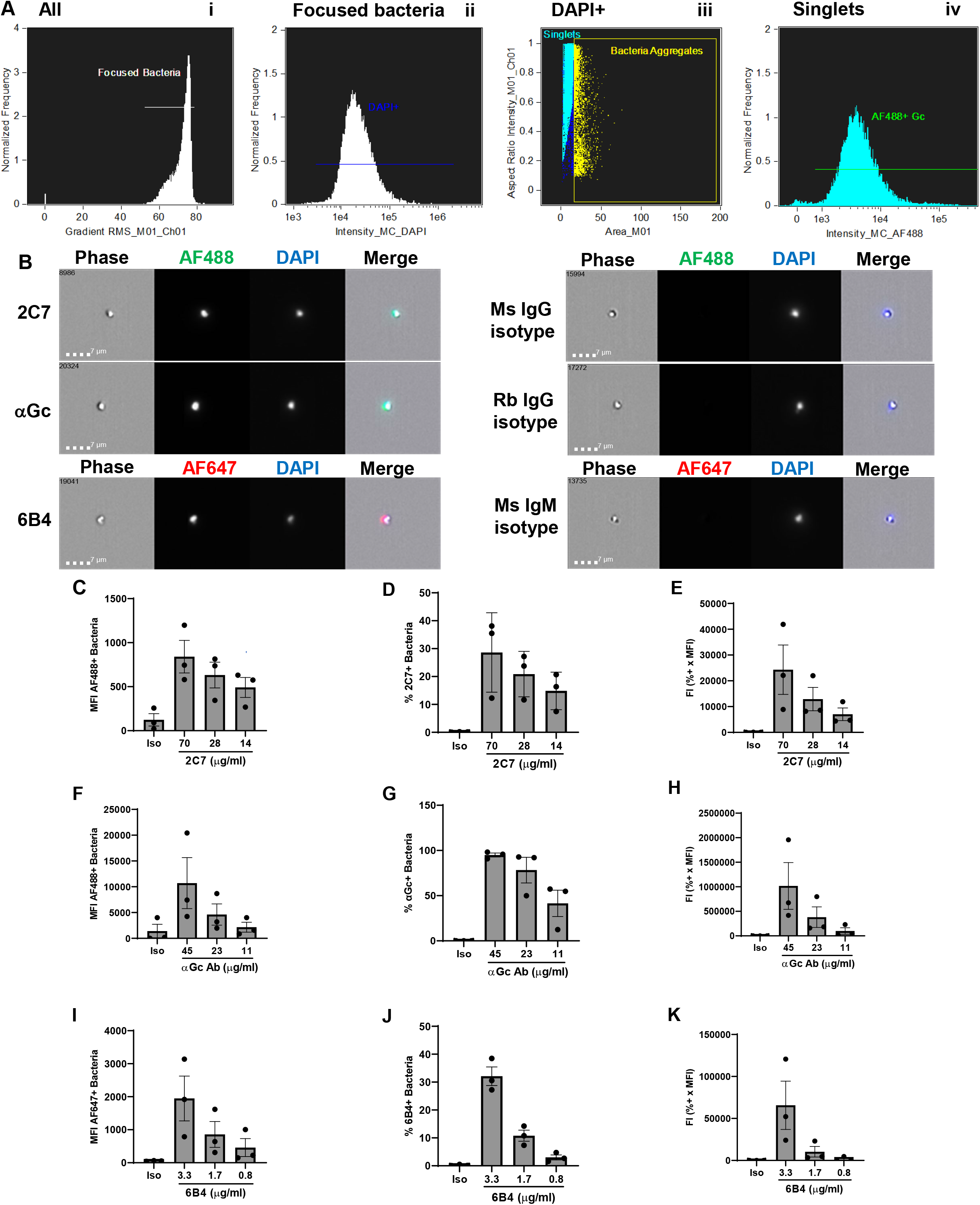
Antibody binding to the surface of *N. gonorrhoeae* by imaging flow cytometry. A) Gating strategy. *N. gonorrhoeae* strain FA1090 was incubated with rabbit anti-*N. gonorrhoeae* antibody and AlexaFluor 488 (AF488)-coupled anti-rabbit antibody, fixed, and incubated with DAPI. On the Imagestream^x^ imaging flow cytometer, samples were gated on (i) RMS of >52, then (ii) focused bacteria, then (iii) DAPI+ singlets, from which AF488-positive fluorescence was measured (iv). B) Representative examples of individual *N. gonorrhoeae* of strain FA1090 that bind mouse (Ms) 2C7 anti-lipooligosaccharide IgG_3_, polyclonal rabbit (Rb) anti-*N. gonorrhoeae* IgG (αGc), and mouse 6B4 anti-lipooligosaccharide IgM (left), but not corresponding isotype controls (right). Number in the upper left corner of each phase image indicates the identity of the particle captured by the imaging flow cytometer. Measurements of mean fluorescence intensity (MFI) (C, F, I), percent positive bacteria (D, G, J), and fluorescence index (FI = MFI x percent positive) (E, H, K) for *N. gonorrhoeae* incubated with the indicated concentrations of 2C7 (C-E), αGc (F-H), and 6B4 (I-K), or highest concentration of each isotype control (iso). Bars indicate the mean ± SD of three independent experiments, with each biological replicate as one data point.

### Serum Bactericidal Activity Assay

*N. gonorrhoeae* was swabbed from GCB agar into liquid media (GCBL) and diluted to an OD_550_ of 0.07. The bacterial suspension was diluted 1:2500 into Hank’s Balanced Salt Solution with Ca^2+^ and Mg^2+^ (HBSS; Gibco, Catalog #14025-092) + 2% BSA to yield ∼ 5 x 10^4^ CFU/ml. Twenty µl of bacterial suspension was added to a 96 well V-bottom plate and incubated with 20 µl of heat-inactivated mouse serum or control antibodies at indicated concentrations, or HBSS + 2% BSA, for 15 min at 37°C, 5% CO_2_. Igd-NHS was added in HBSS + 2% BSA (40 µl) to yield the percent of active complement indicated in **Table 2** for each of the 5 strains used in this study. Samples were mixed with a multichannel pipettor and incubated for 45 min at 37°C, 5% CO_2_. GCBL (100 µl) was added to each sample and mixed vigorously with a multichannel pipettor, and 20 μl of suspension was plated on GCB agar. CFUs were enumerated after overnight incubation at 37°C, 5% CO_2_.

Percent Gc killing was determined by the following equation:

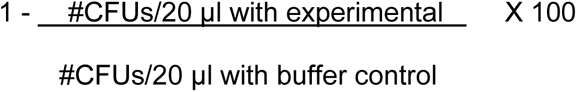

**Table 2.** Serum bactericidal activity elicited by 4CMenB vaccination.

| Strain | SBA Titer of Alum Control* | SBA Titer of 4CMenB* | Normalized SBA Titer% | % NHS# | Positive Control Ab† |
| --- | --- | --- | --- | --- | --- |
| FA1090 | <5 | 40 | > 8 | 2% | 6B4 |
| MS11 | <5 | 160 | > 32 | 0.5% | 6B4 |
| H041 (WHO X) | <5 | 80 | > 16 | 0.5% | 6B4 |
| F62 | <5 | 40 | > 8 | 4% | Rabbit anti-*N. gonorrhoeae* |
| FA19 | 10 | 80 | 8 | 1% | 6B4 |

\* Highest dilution that gave ≥50% killing
% SBA titer of 4CMenB ÷ SBA titer of alum control

Samples with >50% killing at the indicated concentration of antibody or dilution of mouse serum were considered positive.

### PMN Opsonophagocytic Killing Assay

Venous blood was drawn from healthy human subjects who gave informed consent, in accordance with a protocol approved by the University of Virginia Institutional Review Board for Health Sciences Research (#13909). Erythrocytes were removed from heparinized blood by dextran sedimentation, and PMNs were isolated from remaining cells using Ficoll-Hypaque, followed by hypotonic lysis of remaining erythrocytes as described [2]. The remaining cells were > 95% granulocytes by morphology. PMNs were resuspended in Dulbecco’s phosphate buffered saline (D-PBS, without Ca^2+^ and Mg^2+^) plus 0.1% dextrose. PMNs were kept on ice and used within 30 minutes of isolation. Each experimental replicate used PMNs from a different subject.

*N. gonorrhoeae* was swabbed from GCB agar into liquid media (GCBL) and diluted to an OD_550_ of 0.07. The bacterial suspension was diluted 1:2500 into Hank’s Balanced Salt Solution, with Ca^2+^ and Mg^2+^ (HBSS) + 2% BSA to yield ∼ 5 x 10^4^ CFU/ml. Twenty µl of bacterial suspension was added to a 96 well V-bottom plate and incubated with 20 µl of heat-inactivated mouse serum at indicated dilutions or control antibodies for 30 min at 37°C, 5% CO_2_. C6-depleted pooled normal human serum was added (60 µl) to yield a 5% final serum concentration. PMNs (100 µl) at 2 x 10^6^/ml in HBSS + 2% BSA were added, mixed, briefly centrifuged together (600 x g, 4 mins), and incubated for 2 hr at 37°C, 5% CO_2._ Samples were mixed well with a multichannel pipettor, and 20 µl of each suspension was plated onto GCB agar. CFU were enumerated after overnight incubation at 37 °C, 5% CO_2_ and reported as CFU/20 µl, or as percent bacteria killed by the following equation:

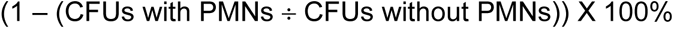

### PMN Opsonophagocytosis of *N. gonorrhoeae* by Imaging Flow Cytometry

Agar-grown FA1090 *N. gonorrhoeae* (1 x 10^8^ CFU) was incubated with 1.5 µl Tag-IT® Violet (TIV) (Biolegend, Catalog # 425101) in 100 µl 1X PBS + 5 mM MgSO_4_. *N. gonorrhoeae* (1 x 10^4^ CFU) diluted in 20 µl of HBSS + 2% BSA was mixed with 20 µl rabbit anti-*N. gonorrhoeae* antibody (Meridian B65111R) diluted 1:50 in of HBSS + 2% BSA (final antibody concentration 50 µg/mL) for 30 minutes. Bacteria were then incubated with 60 µL 5% C6-depleted NHS and 2 x10^6^ primary human PMNs in a total volume of 400 µL to yield a 200:1 PMN:bacterium ratio and centrifuged at 600 x g for 4 min. After 2 hr at 37 °C, 5% CO_2_, cells were fixed without permeabilization, lifted with a cell scraper, blocked in 10% normal goat serum (Gibco, 16210– 064) in PBS, and stained with AF647-labeled goat anti-rabbit antibody to detect extracellular bacteria. Intracellular bacteria were defined as those that were TIV+ and AF647-. Samples were processed for imaging flow cytometry using Imagestream^X^ Mk II with INSPIRE software (Luminex Corporation), and data were analyzed using IDEAS software.

### C3 Deposition on *N. gonorrhoeae*

Strain FA1090 bacteria was prepared as above, with 25 µl of 2C7 (1:2 dilution, 0.14 mg/ml) added as the opsonic antibody for 20 min at 37°C, 5% CO_2_. After 2 washes with RPMI + 10% HI-FBS, C6-depleted NHS (in RPMI + 2% BSA) was added to the indicated final concentration and incubated for 15 mins at 37°C, 5% CO_2_. Bacteria were washed, then incubated with FITC-conjugated goat IgG anti-human C3 (1:5000 dilution) for 15 min at 37 °C, 5% CO_2_. Bacteria were washed once with PBS and resuspended in 2% PFA containing 5 µg/ml DAPI. Samples were analyzed by imaging flow cytometry using ImagestreamX Mk II with INSPIRE® software. FITC fluorescence was detected with excitation at 488 nm and emission collected with a 480– 560 nm filter. DAPI fluorescence was detected as above. Single color samples for FITC and DAPI were collected without Brightfield to create a compensation matrix for each experiment. Gating strategy was as for antibody binding to *N. gonorrhoeae*.

### HL-60 Opsonophagocytosis Assay

HL-60 cells (ATCC Catalog # CCL-240) were maintained in Iscove’s Modified Dulbecco’s Medium (IMDM, Corning Catalog #15-016-CV) with 20% heat-inactivated fetal bovine serum (FBS; Hyclone, Catalog # SH30071.03), Glutamax (Gibco, Catalog #35050-061) and antibiotic-antimycotic (Gibco, Catalog #15240-062). Cells were split to a density of 1.5 x 10^5^ /ml and not allowed to exceed 1 x 10^6^/ml. For differentiation, cells were centrifuged in 15 ml conical tubes at 300 x g for 7 min, then resuspended at 1 x 10^6^/ml in growth medium containing 1.25% dimethyl sulfoxide (DMSO, Sigma, Catalog #D2650) for 3 days at 37 °C, 5% CO_2_. Cells were then changed into growth medium without DMSO for an additional 2 days at 37° C, 5% CO_2_. Cells were washed 3 x in RPMI and resuspended at 2 x 10^7^/ml in RPMI + 2% BSA. Under these differentiation conditions, HL-60 were >90% viable and >70% CD11b+ by flow cytometry.

*N. gonorrhoeae* was labeled with Tag-IT Violet as above, then washed and resuspended at 1.6 x 10^9^ CFU. Bacteria (12.5 µl for 2 x 10^6^/well) were added to a V-bottom 96 well plate. 2C7 IgG (12.5 µl, 0.28 mg/ml) was added and the plate incubated for 30 min at 37°C, 5% CO_2_. Samples were washed 3 times before C6-depleted NHS (50 µl of 4% for 1% final) and differentiated HL-60 cells (100 µl of 2 x 10^7^/ml) were added and incubated for 1 hr at 37°C, in 5% CO_2._ Samples were washed 1 time then Zombie Green (Biolegend, Catalog #423111, 1:1000) was added to measure and gate on the % live HL-60 cells and anti-mouse/human CD11b PE (Biolegend, Catalog #101207, 1:50) was added to measure and gate on differentiated HL-60 cells. Samples were incubated for 30 min on ice, washed 1 x in PBS and fixed with 2% PFA.

Analysis of 1 x 10^4^ cells per sample was performed by imaging flow cytometry using ImagestreamX Mk II with INSPIRE® software (Luminex Corporation). A scatter plot with Area_M01 on the X axis and Aspect Ratio_M01 on the Y axis was used to identify single HL-60 cells with Area_M01 between 200 and 400 and Aspect Ratio_M01 between 0.6 and 1. From the singlets gate, focused cells were identified with Brightfield Gradient Root Mean Square (RMS) values over 50. Dead cells with Zombie Green intensity over 1e4 were excluded. From the live cell population, differentiated HL-60s were defined as cells positive for CD11b PE. Using a non-differentiated control, an intensity over 5e3 was considered CD11b+. Tag-it Violet® fluorescence was measured in the live, differentiated cell population. Cells with TIV intensity over 6e3 were considered associated with Gc. Zombie Green fluorescence was detected with excitation at 488 nm and emission collected with a 480-560 nm filter. PE fluorescence was detected with excitation at 561 nm and emission collected with a 560-595 nm filter. Tag-IT Violet® fluorescence was detected with excitation at 405 nm and emission collected with a 420-505 nm filter. Single color samples for Zombie Green, PE, and Tag-it Violet® were collected without Brightfield to create a compensation matrix for each experiment.

### Statistics

Independent biological replicates were conducted with different bacterial cultures on different days, different preparations of HL-60 cells, and/or different human subjects’ PMNs. Statistical comparisons were made as indicated in the figure legends.

## Results

### Antibody binding to the surface of *N. gonorrhoeae*

Imaging flow cytometry was applied to assess the binding of antibodies to antigens on the surface of live *N. gonorrhoeae,* with the goal to use immunized serum as the antibody source. Imaging flow cytometry has a slower flow rate than conventional flow cytometry and can better distinguish small particles like individual *N. gonorrhoeae* (diameter ∼0.5 µm) from bacterial aggregates, whose fluorescence could skew results. Our group has previously used imaging flow cytometry to measure binding of the N-terminal domain of human carcinoembryonic antigen-related cell adhesion receptors to opacity protein adhesins on *N. gonorrhoeae* (55), binding of complement iC3b to fluorescent beads (56), and binding of complement C4b-binding protein to *N. gonorrhoeae* (57). As proof of concept, live *N. gonorrhoeae* strain FA1090 was incubated with one of three purified antibodies: 2C7, a mouse monoclonal IgG_3_ antibody that recognizes an epitope including the β chain lactose of *Neisseria* lipooligosaccharide (58, 59); a polyclonal rabbit IgG raised against *N. gonorrhoeae* (60); and 6B4, a mouse monoclonal IgM antibody that recognizes the lacto-*N-* tetraose α chain of *Neisseria* lipooligosaccharide (58, 61). Bacteria were washed, incubated with an Alexa Fluor 488® (AF488)-IgG or Alexa Fluor 647® (AF647)-IgM coupled secondary antibodies, resuspended in fixative, counterstained with DAPI, and analyzed by imaging flow cytometry. Primary antibodies of the same isotype with the same secondary antibodies were used as negative controls for background fluorescence. The gating strategy is depicted in **Fig 1A**: (i) Focused particles were identified as those with high gradient Root Mean Square (RMS), which is a measure of image sharpness based on pixelation. (ii) Within this gate, DAPI+ particles (*N. gonorrhoeae*) were identified. (iii) DAPI+ bacterial aggregates, as defined by their area and aspect ratio, were excluded from further analysis. (iv) The intensity of AF488 IgG+ DAPI+ singlets was quantified. Examples of antibody-positive bacteria are presented on the left of **Fig 1B**, and matching isotype controls on the right.

Both 2C7 and the anti-*Neisseria gonorrhoeae* polyclonal rabbit IgG elicited a concentration-dependent increase in AF488 fluorescence of *N. gonorrhoeae*, while there was no-to-marginal detection of fluorescence of the isotype control in each condition **(Fig 1C-H).** The mouse IgM that recognizes the lacto-*N-*tetraose α chain of *Neisseria* lipooligosaccharide, 6B4, also elicited a concentration-dependent increase in AF647 fluorescence of *N. gonorrhoeae* **(Fig 1 I-K).** The magnitude of fluorescence was calculated in three ways: 1) mean fluorescence intensity (MFI) **(Fig C, F, I)**; 2) percent positive bacteria (e.g. in the green gate in **Fig 1A iv**) **(Fig 1D, G, J)**; 3) fluorescence index, calculated as the MFI multiplied by the percent positive bacteria **(Fig 1E, H, K)**. The use of hybridoma supernatants of 2C7 and 6B4 precluded us from incubating *N. gonorrhoeae* with higher concentrations of antibody; hence the percent positive bacteria in **Fig 1D** and **Fig 1J** do not reach saturation. However, these lower concentrations may be more reflective of specific antibodies in serum from immunized animals. Thus (imaging) flow cytometry can be used to measure the binding of antibodies to the surface of *N. gonorrhoeae,* a prerequisite for functional antibody activities.

### Serum bactericidal activity assay for *N. gonorrhoeae*

SBA is a defined correlate of protection for vaccination against extracellular Gram-negative bacteria, including the related *N. meningitidis* (38, 39, 62–64). We sought to develop a SBA platform with *N. gonorrhoeae* to allow for screening of serum from immunized and control animals in a rapid and reproducible manner. To establish this platform, we used 6B4 IgM antibody, since pentameric IgM is a potent initiator of the classical complement cascade. Ig-depleted normal human serum (Igd-NHS) was then added as the source of complement. HBSS served as the medium for incubation. The use of Igd-NHS removes any potential contribution of naturally occurring antibodies in human serum that cross-react with *N. gonorrhoeae.* CFU were enumerated after overnight growth.

Using strain FA1090, there was an antibody- and complement-dependent effect on *N. gonorrhoeae* viability (**Fig 2A**). As expected for this serum-resistant strain, there was no effect of NHS on bacterial viability in the absence of 6B4 (black line, **Fig 2A**). At each concentration of NHS, increasing concentrations of 6B4 led to reduced bacterial survival. At the highest concentrations of 6B4, few to no CFU were enumerated (red, orange lines, **Fig 2A**); there was a concentration-dependent decrease in bacterial killing as the concentration of antibody decreased (**Fig 2A**). A concentration of 2% NHS and 6B4 hybridoma supernatant at 0.2 µg/ml yielded approximately 50% bacterial survival compared with the HBSS control (light blue vs. black symbols, **Fig 2A**). We selected 2% NHS as the concentration of complement source that yielded the greatest dynamic range of SBA against strain FA1090. Similar calculations were made for other strains of *N. gonorrhoeae* (**Table 2**).

**Figure 2.**
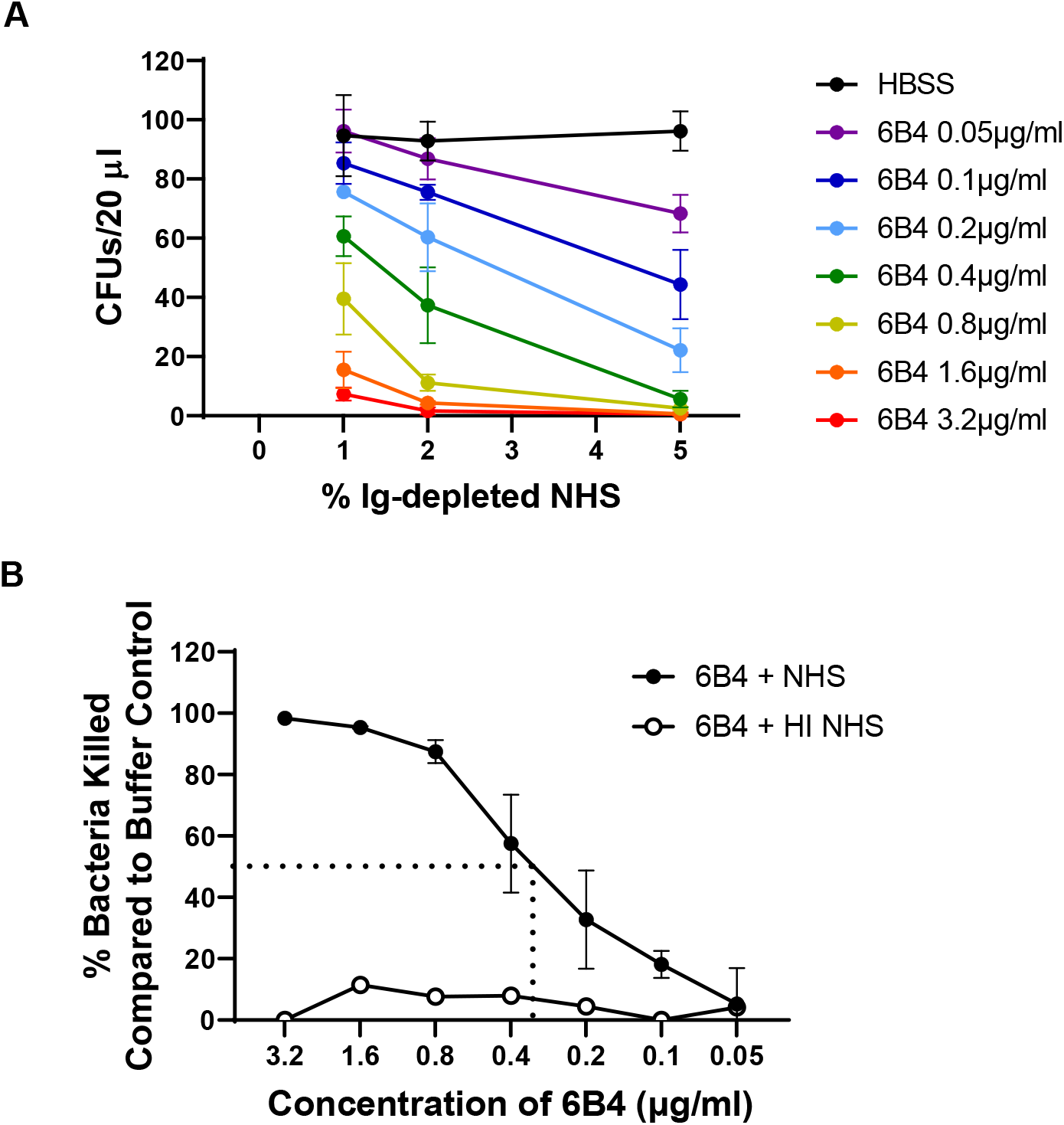
*N. gonorrhoeae* serum bactericidal assay. A) *N. gonorrhoeae* strain FA1090 (10^3^ CFU) was incubated with the indicated concentration of 6B4 or HBSS buffer, followed by addition of the indicated concentration of Ig-depleted normal human serum (NHS). Bacterial colony forming units (CFU) from 20 µl of the incubation medium were enumerated after overnight growth on GCB agar. B) Bacteria killed were expressed as a percent of the buffer (HBSS) control and graphed relative to the concentration of 6B4 in 2% NHS (solid circles). The dotted lines indicate the extrapolation to the 6B4 concentration that yields 50% bacterial killing. The SBA titer, or the lowest concentration that yielded ≥ 50% killing, was calculated as 0.4 µg/ml (red). 5% heat-inactivated (HI) NHS, the highest concentration of NHS that was tested, served as a negative control (open circles). Bars indicate the mean ± SD of three independent experiments, with each biological replicate as one data point.

To calculate the SBA titer, strain FA1090 was incubated with increasing dilutions of 6B4 prior to exposure to 2% NHS. **Fig 2B** reports the percent of *N. gonorrhoeae* killed, relative to bacteria incubated with the buffer control, for the indicated concentrations of 6B4. The SBA titer was calculated as the lowest concentration of antibody in which ≥50% of the bacteria were killed (dotted line showing interpolation for 50%; SBA titer = 0.4 µg/ml, **Fig 2B**). Of note, there was no appreciable killing of *N. gonorrhoeae* when heat-inactivated (HI) NHS was provided (white symbols, **Fig 2B**), indicating that bactericidal activity required active complement. A similar approach was used to establish the SBA titer for four other strains of *N. gonorrhoeae* commonly used for biomedical experimentation and vaccine development: MS11, H041 (WHO X), F62, and FA19. The concentration of Ig-depleted NHS and the positive control antibody for each of these strains is presented in **Table 2**. Of note, we measured positive SBA titers not only for the serum-sensitive strains F62, MS11, and H041, but also the two highly serum-resistant strains FA1090 and FA19.

For evaluating immunized and control mouse serum, our plan for measuring and reporting SBA is as follows: 1) Determine the SBA titer for the serum from control mice. 2) Measure bacterial killing by the immunized serum, starting with 2x the dilution of the SBA titer for the serum from control mice. 3) If >50% of the bacteria are killed by the immunized serum at this dilution, conduct SBA over a full range of dilutions of serum and calculate the SBA titer. 4) Report the results as a normalized SBA titer. If there is no increased SBA with the immunized serum, report as a normalized SBA titer of 2.

### Antibody-elicited opsonophagocytic killing activity against *N. gonorrhoeae*

Gonorrhea is typified by influx of neutrophils at the site of infection (14), and opsonophagocytic killing may contribute to the efficacy of *N. meningitidis* vaccines (41, 65, 66). To assess the contribution of immunization-elicited antibodies in opsonphagocytic killing of *N. gonorrhoeae,* we established an *in vitro* assay for opsonphagocytic killing activity (OPKA) using primary human PMNs from the venous blood of healthy human subjects. As proof of concept, *N. gonorrhoeae* was first incubated with an anti-*N. gonorrhoeae* rabbit polyclonal IgG (see **Fig 1**), followed by addition of C6-depleted NHS as a complement source. PMNs were then added at a ratio of 200 per bacterium, and after 2 hr of coincubation, CFU were enumerated. C6-depleted serum, which cannot make the membrane attack complex and cannot elicit SBA, was used to allow the effect of PMN-dependent killing to be selectively assessed. Binding and phagocytosis of *N. gonorrhoeae* by PMNs under the OPKA conditions were examined by imaging flow cytometry, using bacteria prelabeled with the fluorophore Tag-IT Violet. After incubation, PMNs were fixed but not permeabilized, and an Alexa Fluor 647-coupled antibody was used to detect extracellular bacteria. As shown in the representative images in **Fig 3A**, *N. gonorrhoeae* was phagocytosed by PMNs (Tag-IT Violet+, Alexa Fluor 647-). At the MOI used, approximately 4% of all PMNs had associated *N. gonorrhoeae,* and 1.8% of all PMNs had phagocytosed bacteria.

**Figure 3.**
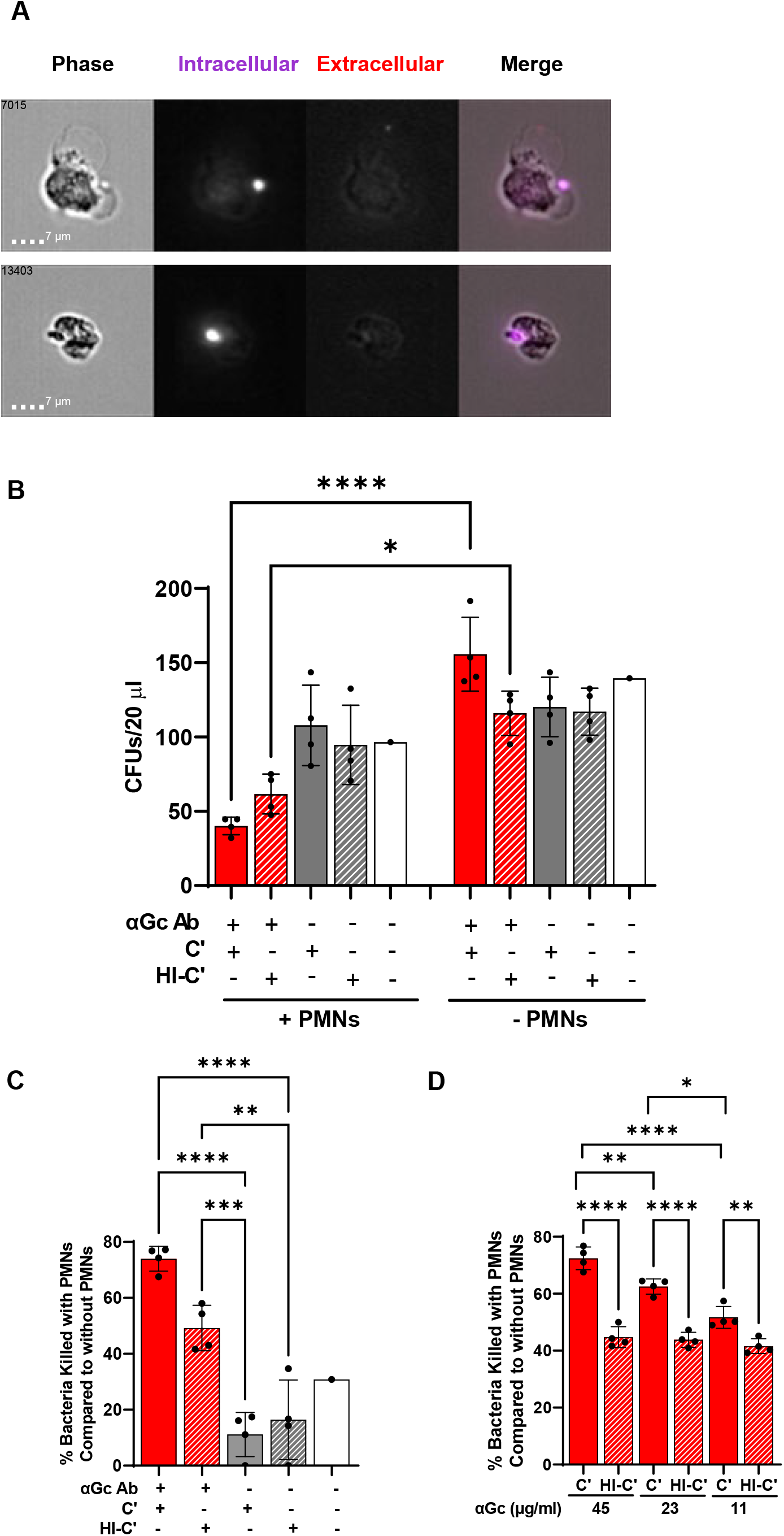
Opsonophagocytic killing assay with primary human PMNs. A) Examples of two PMNs that underwent opsonophagocytosis of Tag-IT Violet-labeled (purple) *N. gonorrhoeae* strain FA1090, in the presence of C6-depleted NHS and rabbit anti-*N. gonorrhoeae* antibody (50 µg/ml), analyzed by imaging flow cytometry. PMNs were fixed and stained with Alexa Fluor 647-coupled anti-*N. gonorrhoeae* antibody to detect bound but not phagocytosed bacteria (red). Number in the upper left corner of each phase image indicates the identity of the cell captured by the imaging flow cytometer. (B) *N. gonorrhoeae* strain FA1090 was mixed with rabbit anti-*N. gonorrhoeae* antibody or buffer control as above, except not fluorescently labeled. C6-depleted normal human serum that was untreated (C’) or heat-inactivated (HI-C’) was added to 5% final concentration, along with primary human PMNs (+PMNs) or buffer (-PMNs). CFU were enumerated from 20 µl of the incubation mix. Red solid bar with PMNs is the full opsonophagocytic condition. C) Results are as in B except each condition was expressed as a percentage of the enumerated CFU for the same condition in the absence of PMNs. D) Results are as in B except with the indicated concentrations of rabbit anti-*N. gonorrhoeae* antibody, and presented as the percent of bacteria killed compared to bacterial killing when heat-inactivated serum was used. In B-D, results presented are the average ± SD from 4 independent experiments using different human subjects’ PMNs, with each biological replicate as one data point. *p≤0.05, **p≤0.005, ***p≤0.001, ****p≤0.0001 by one-way ANOVA followed by Holm-Šidák multiple comparisons test.

For OPKA, CFU were enumerated for strain FA1090 *N. gonorrhoeae* with or without antibody, in the presence of intact or heat-inactivated C6-depleted NHS, in the presence or absence of PMNs from three unrelated subjects. Addition of antibody significantly reduced the viability of *N. gonorrhoeae* exposed to PMNs compared to those that were not, whether active complement or heat-inactivated complement was used (compare solid red bars with and without PMNs, and red hatched bars with and without PMNs, **Fig 3B**). Reductions in bacterial viability in the presence of PMNs were not significant in other conditions **(Fig 3B)**. These observations agree with a previous report on the effect of IgG on phagocytic killing of *N. gonorrhoeae* (60).

To take into account any differences in viability that occurred independently of PMNs (right side, **Fig 3B**), CFU in the experimental condition was divided by CFU from the same conditions without PMNs, subtracted from 1 to convert into fraction of bacteria killed, and multiplied by 100%. From these results, we measured a statistically significant decrease in *N. gonorrhoeae* viability when antibody, complement, and PMNs were all present, relative to antibody alone, active complement alone, or neither **(Fig 3C).** The effect of antibody was concentration-dependent, with statistically significant increased bacterial survival with increasing dilutions of the opsonic antibody **(Fig 3D).** However, this was dependent on active complement, because there was no difference in the magnitude of bacteria killed by PMNs at these different antibody concentrations when heat-inactivated serum was used as complement source **(Fig 3D).** We conclude that *N. gonorrhoeae-*specific IgG in the presence of active complement and PMNs elicits OPKA. However, the magnitude of this response is limited, compared with bacteria like pneumococci that are strongly controlled by vaccine-elicited OPKA (67). These results align with studies demonstrating the myriad ways *N. gonorrhoeae* resists killing by PMNs (14).

### Alternative assays for examining functional antibody contributions to control of *N. gonorrhoeae*

The premise of the SBA and OPKA assays is that antibodies facilitate complement deposition on *N. gonorrhoeae,* leading to membrane attack complex formation and opsonophagocytosis, respectively. An alternative readout of these activities is deposition of complement component 3 (C3) on the bacterial surface, which occurs downstream of the classical complement cascade. We evaluated the ability of the monoclonal anti-LOS antibody 2C7 and C6-depleted NHS to elicit C3 deposition on strain FA1090 *N. gonorrhoeae* using imaging flow cytometry, since 2C7 has been previously reported to increase C3 on the surface of *N. gonorrhoeae* (59, 68). C6-depleted NHS was used to avoid bacterial lysis, which could confound quantitation of bacterial C3 positivity. Bacteria were mixed with 2C7 antibody or isotype control, after which C6-depleted NHS at the indicated concentrations or buffer were added, following the initial steps outlined above for OPKA. Bacteria were then fixed and stained with a fluorescent antibody that recognizes all forms of human C3, including the opsonins C3b and iC3b. The intensity of C3 binding was calculated as the fluorescence index (percent positive x MFI). C3 deposition was only detected in the presence of C6-depleted NHS, and was markedly increased when 2C7 was added, compared with isotype control (**Fig 4A**). This assay can be used to demonstrate increased complement deposition on *N. gonorrhoeae* that is promoted by vaccine-elicited antibodies.

**Figure 4.**
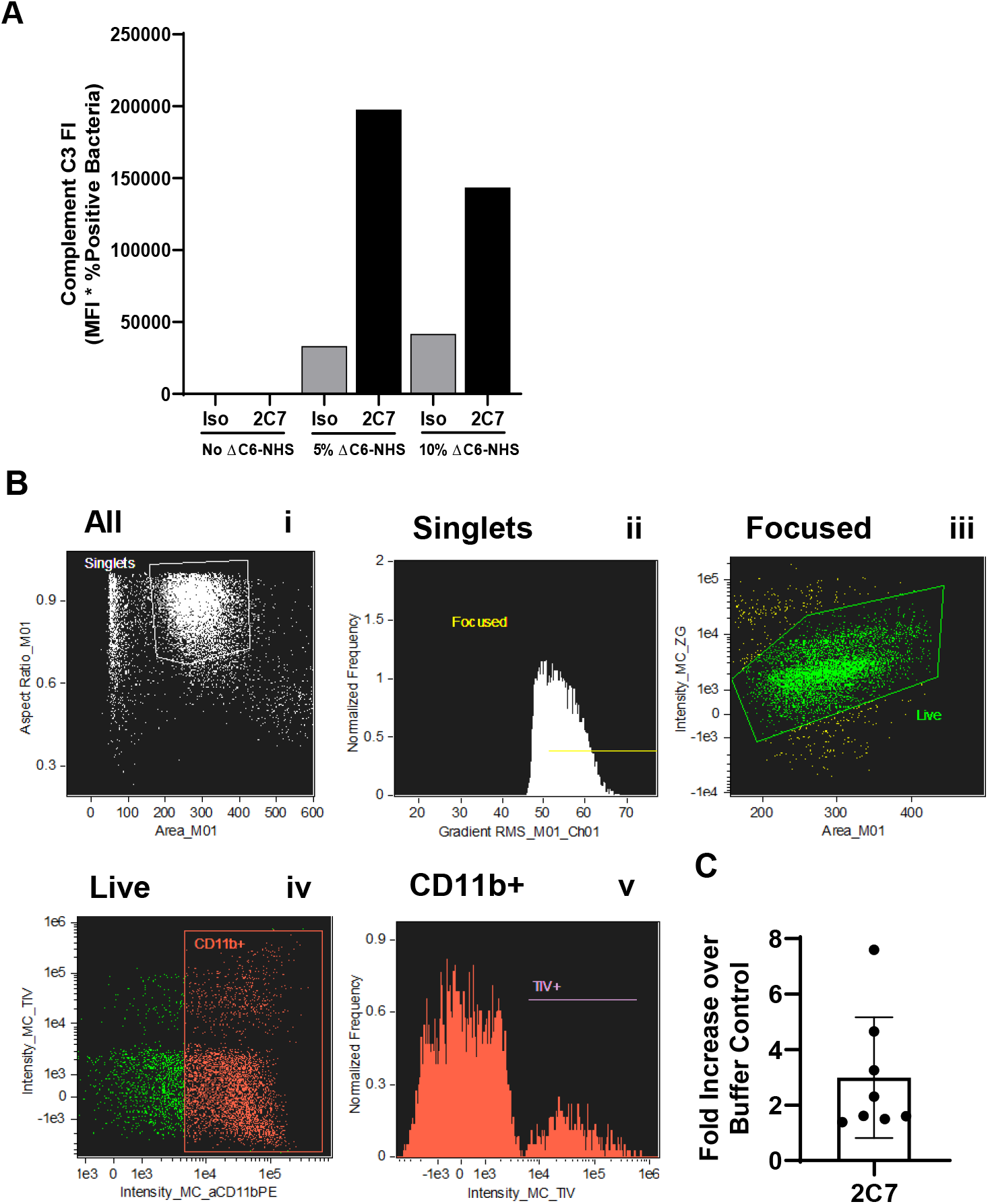
Alternative assays for complement-mediated functional antibody activity against *N. gonorrhoeae.* A) Complement C3 deposition. *N. gonorrhoeae* strain FA1090 was incubated with 2C7 or HBSS in the absence or presence of the indicated concentrations of C6-depleted NHS. Bacteria were fixed, stained with a FITC-labeled anti-human C3 antibody, counterstained with DAPI, and processed for flow cytometry. The fluorescence index (FI) was calculated as in Fig 1. One representative experiment is reported. B-C) Opsonophagocytic assay with differentiated HL-60 cells. *N. gonorrhoeae* strain FA1090 was labeled with Tag-IT Violet (TIV), then incubated with 2C7 or isotype control, followed by exposure to C6-depleted NHS and differentiated HL-60 cells. After 2 hr, cells were fixed, stained with anti-CD11b for differentiation and Zombie Green for viability, and processed for imaging flow cytometry. Gating strategy: i) Singlets with size and aspect ratio indicative of intact cells; ii) Focused singlets; iii) Live cells; iv) CD11b+ cells. In v), the fluorescence intensity of TIV+ *N. gonorrhoeae* was measured in the CD11b+ gate. C) The fold increase in opsonophagocytosis of *N. gonorrhoeae* by dHL-60 cells elicited by 2C7 compared with the buffer control is reported. Bars represent the mean ± SD of 8 independent experiments conducted on different days, with each biological replicate as one data point.

Experiments with primary PMNs require human subjects approval and have the potential for person-to-person variability. As an alternative to human blood cells, we used the human promyelocytic cell line HL-60, differentiated towards a neutrophil-like phenotype by treatment with DMSO (dHL-60). HL-60 cells have been used extensively for assaying OPKA against *S. pneumoniae* and other Gram-positive bacteria (69, 70). However, in our hands dHL-60 cells do not kill *N. gonorrhoeae* (56). As a surrogate, we established an imaging flow cytometry assay using Tag-IT Violet-labeled *N. gonorrhoeae,* where we hypothesized the intensity of Tag-IT Violet fluorescence of dHL-60 would be proportional to the extent of phagocytosis. dHL-60 cells were incubated with *N. gonorrhoeae,* IgG antibody against *N. gonorrhoeae* (in this case, 2C7), and C6-depleted NHS. Cells were then fixed and stained with dyes to assess cellular viability (Zombie) and differentiation (CD11b upregulation). The percent of dHL-60 cells that were Tag-IT Violet+ and the mean fluorescence intensity for Tag-IT Violet+ cells was determined from the viable, CD11b+ focused singlet population (**Fig 4Bv**). While 2C7 elicited increased opsonophagocytosis compared with the buffer (no antibody or complement) control, the spread of the biological replicates was greater than expected for a cell line, and greater than observed for OPKA with human PMNs (**Fig 4C**; compare with data points on solid red bar, **Fig 3C**). Given the expenses of cell culture and flow cytometry, the inconsistent dHL-60 response, and ability of primary human PMNs to kill *N. gonorrhoeae* in a complement- and antibody-dependent manner, our preference is to use human PMNs for evaluating antibodies elicited by immunization.

### The 4CMenB meningococcal serogroup B vaccine elicits functional antibody responses against multiple strains of *N. gonorrhoeae*

The reverse vaccinology-engineered meningococcal serogroup B vaccine 4CMenB has been shown to elicit antibodies in humans and mice that recognize *N. gonorrhoeae* and protect mice from gonococcal infection (26, 27, 29). To examine the ability of 4CMenB to promote functional antibody responses against *N. gonorrhoeae,* mice were vaccinated with 4CMenB or alum as a control. Serum was collected on day 63 post-immunization, pooled, heat-treated to inactivate mouse complement, and assessed for antibody binding, SBA, and OPKA, using the human sera described above as the complement sources.

#### Antibody binding

Five different strains of *N. gonorrhoeae* were assessed for 4CMenB elicited antibody binding to the bacterial surface, reported as fluorescence index. Here, *N. gonorrhoeae* was mixed with serum from 4CMenB-vaccinated mice, or alum-treated mice as the control, prior to the staining procedure described for **Fig 1**. Purified antibody that recognizes the given strain of *N. gonorrhoeae* served as positive control (see **Table 2**), and isotype antibody as negative control. For the tested strains except H041, the fluorescence index of IgG bound to *N. gonorrhoeae* was markedly increased when using serum from 4CMenB-vaccinated mice, compared with alum-treated mice or the isotype control **(Fig 5).** Among the four strains with increased binding with 4CMenB, there were differences in both the fluorescence index and the magnitude of the difference between 4CMenB-vaccinated and alum-treated mice, with strain F62 having the most robust binding of IgG from 4CMenB, and strains FA19 and MS11 the least (F62>H041>FA1090>FA19∼MS11). These differences could help inform future studies to identify the gonococcal antigens cross-reacting with 4CMenB using comparative proteomic analyses (33).

**Figure 5.**
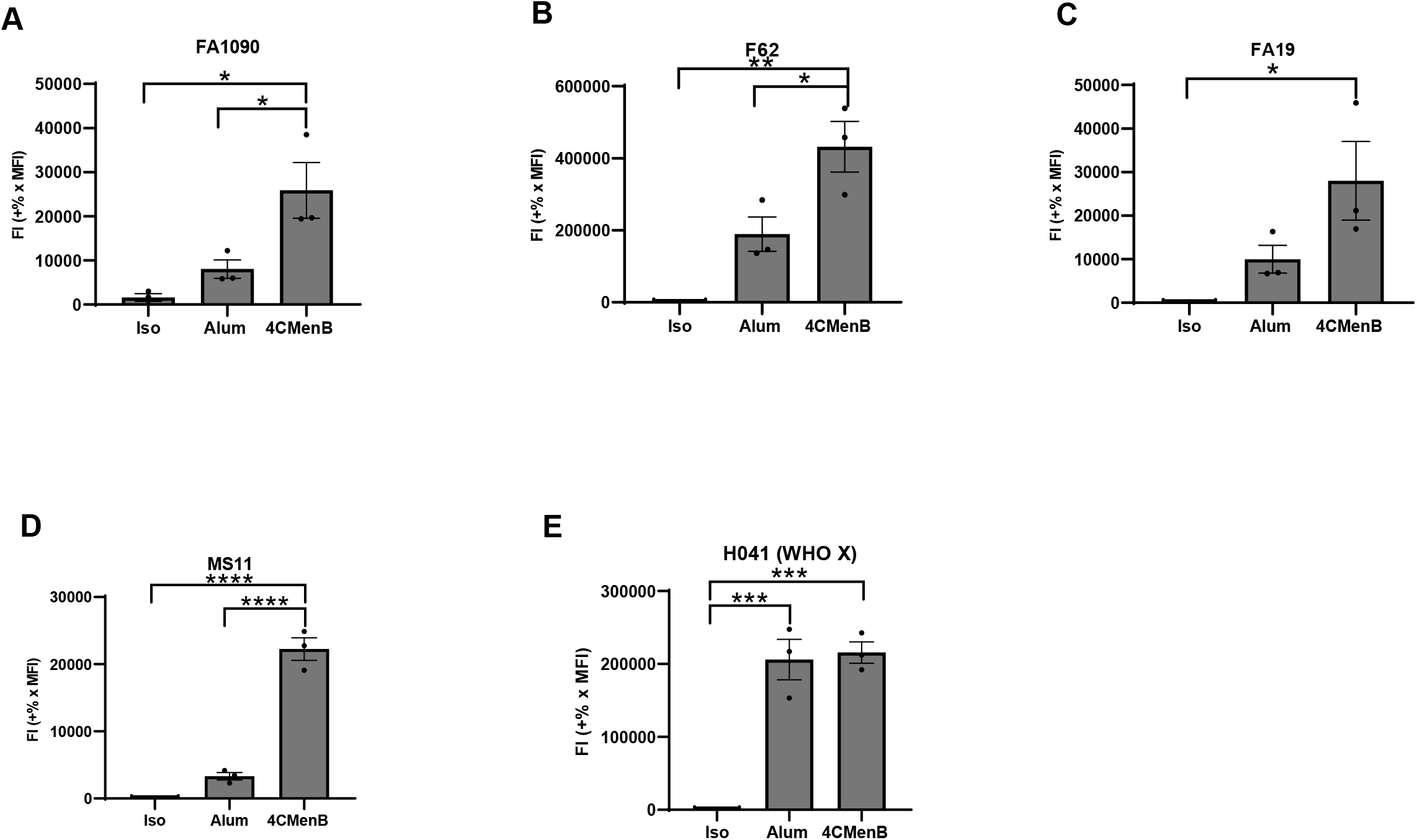
4CMenB vaccination of mice elicits cross-reactive antibodies that bind the surface of multiple strains of *N. gonorrhoeae.* Bacteria of the indicated strain background were incubated with a 1:10 dilution of pooled serum from 4CMenB-vaccinated or alum treated mice, or mouse IgG isotype control. Bacteria were fixed, processed for imaging flow cytometry, and fluorescence index calculated as in Fig 1. Bars indicate the mean ± SD of three independent experiments, with each biological replicate as one data point. *p≤0.05, **p≤0.005, ***p≤0.001, ****p≤0.0001 by ordinary one-way ANOVA with Tukey’s Multiple Comparison’s test.

#### Serum bactericidal activity

Strain FA1090 was subjected to SBA as in **Fig 2**, using pooled serum from 4CMenB-vaccinated and alum-treated mice. Unlike purified antibodies, we found that serum from the alum-treated mice killed *N. gonorrhoeae* in the presence of active complement (not shown). We hypothesized that the presence of nonspecific or low-affinity antibodies were responsible for SBA elicited by the unimmunized controls. To mitigate this effect, BSA at concentrations ranging from 0.5-5% was added to the assay buffer (HBSS). While 0.5% BSA did not prevent killing of the strain FA1090 by the serum from alum-treated mice at dilutions up to 1:10 (red bars, **Fig 6A**), addition of 2% BSA reduced the SBA background of alum-treated mouse serum (blue bars, **Fig 6A**) without impeding the SBA elicited by the serum from 4CMenB-vaccinated mice (blue bars, **Fig 6B**). 5% BSA further reduced the background of the serum from alum-treated mice, but also reduced SBA elicited by the 4CMenB-vaccinated serum (green bars, **Fig 6A-B**). These effects were specific to the presence of serum, since addition of BSA did not affect the SBA of the two positive control antibodies, 6B4 **(Fig 6C)** and rabbit IgG anti-*N. gonorrhoeae* **(Fig 6D).** Based on these results, going forward, HBSS + 2% BSA was used as the diluent for SBA assays.

**Figure 6.**
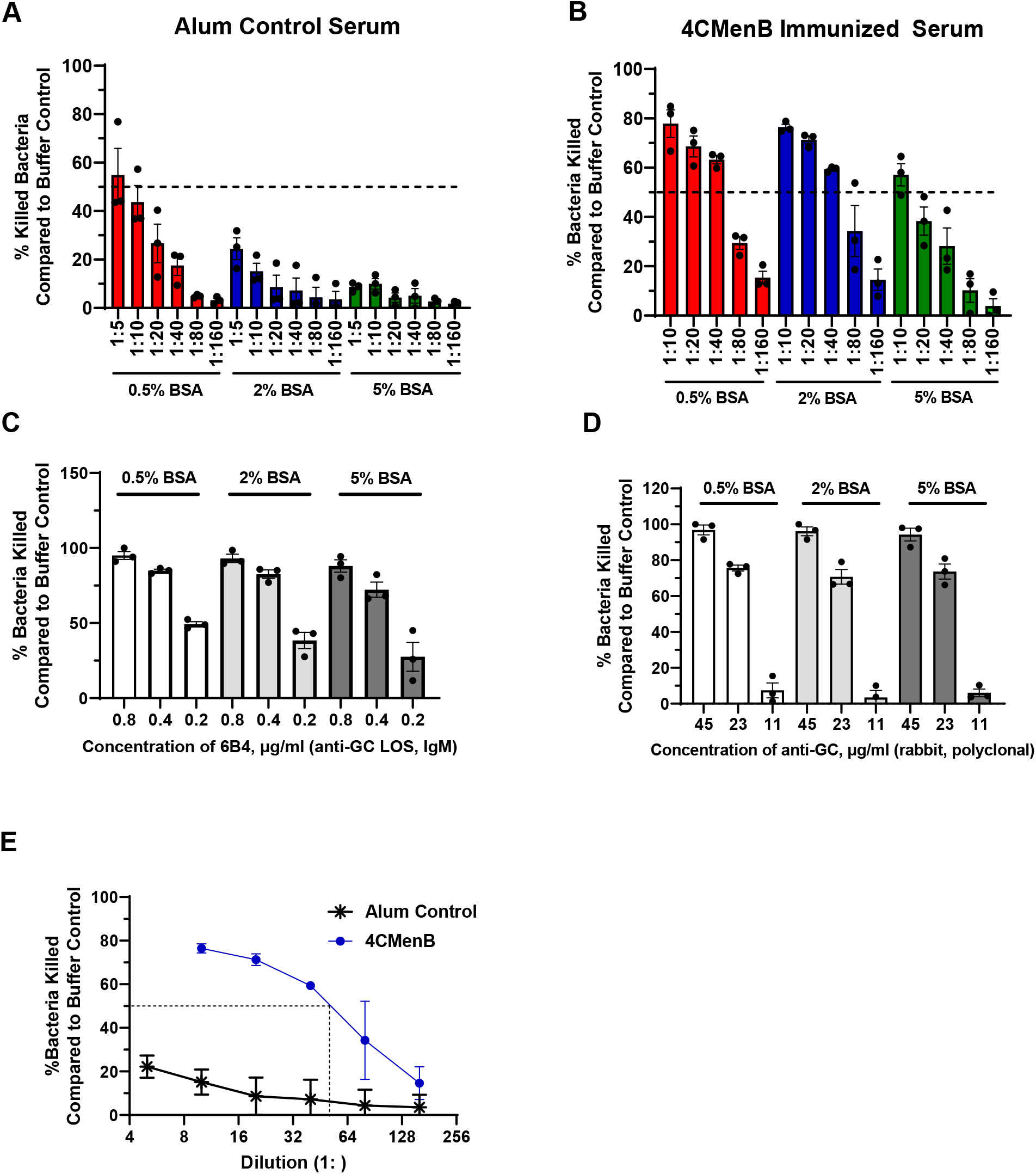
Serum bactericidal activity elicited by 4CMenB vaccination against *N. gonorrhoeae.* (A-B) *N. gonorrhoeae* strain FA1090 was incubated with the indicated concentrations of BSA in the presence of the indicated dilution of pooled serum from alum treated (A) or 4CMenB-vaccinated (B) mice. Serum bactericidal activity was measured using 2% Ig-depleted NHS as complement source and reported as percent bacteria killed relative to the buffer control. (C-D) *N. gonorrhoeae* strain FA1090 was mixed with the indicated concentration of BSA and 6B4 (C) or rabbit anti-*N. gonorrhoeae* (Gc) antibody (D). The percentage of bacteria killed relative to buffer control was calculated as in A-B. Based on the results, 2% Ig-depleted NHS, 2% BSA, and 0.4 µg/ml 6B4 or 22.5 µg/ml rabbit anti-*N. gonorrhoeae* antibody were selected as positive control conditions for FA1090 *N. gonorrhoeae;* similar calculations were made for other *N. gonorrhoeae* strains with the positive control antibodies in Table 2. (E) SBA activity measured using strain FA1090 *N. gonorrhoeae,* 2% BSA, the indicated dilutions of pooled serum from 4CMenB-vaccinated (circle) or alum-treated control (asterisk) mice, and 2% Ig-depleted NHS. SBA titers for all five strains of *N. gonorrhoeae* are reported in Table 2. Bars indicate the mean ± SD of three independent experiments, with each biological replicate as one data point.

We next determined the SBA titer of 4CMenB-vaccinated serum against strain FA1090 and 4 other strains of *N. gonorrhoeae* using 2% BSA in the assay. The SBA response was more potent than the alum control in each case, with a calculated SBA titer of 40 for 4CMenB against FA1090 compared with <4 for alum (**Fig 6E**). SBA titers for 4CMenB ranged from 40-160 against MS11, H041, F62 and FA19 compared to <5 for their respective alum-controls (except for FA19, whose alum SBA titer was 10) (**Table 2**). The normalized SBA titer, or the SBA titer elicited by the vaccinated serum (*e.g.* 4CMenB) divided by the SBA titer elicited by the control serum (*e. g.* alum treated), is also presented in **Table 2** for each strain. These findings show that 4CMenB vaccination elicits a humoral response that promotes SBA against multiple strains of *N. gonorrhoeae*.

#### Opsonophagocytic killing assay

We applied the OPKA assay with primary human PMNs to strain FA1090 *N. gonorrhoeae* that was incubated with serum from mice vaccinated with 4CMenB, or treated with alum. The numbers of PMNs and the volume of mouse serum that would be needed precluded us from calculating an OPKA titer. Instead, we selected a single dilution (1:10) of vaccine-elicited or control serum for OPKA measurements and expressed vaccine-elicited OPKA as a fold-increase over its biological control. As in the SBA, 2% BSA was added to *N. gonorrhoeae* prior to adding the mouse serum.

Exposure to C6-depleted NHS, serum from 4CMenB-vaccinated mice, and human PMNs reduced survival of *N. gonorrhoeae* relative to all other conditions (blue bar on left, **Fig 7A**). The serum from alum-treated mice had no effect on OPKA compared with bacteria that were not incubated with any mouse serum (red bar vs. green bar on left, **Fig 7A**). In the absence of PMNs, there was no reduction in bacterial CFU (right half of **Fig 7A**). In fact, non-heat-inactivated mouse serum supported *N. gonorrhoeae* outgrowth, potentially due to the presence of heat-sensitive nutrients that *N. gonorrhoeae* can consume (red and blue bars, right half of **Fig 7A**).

**Figure 7.**
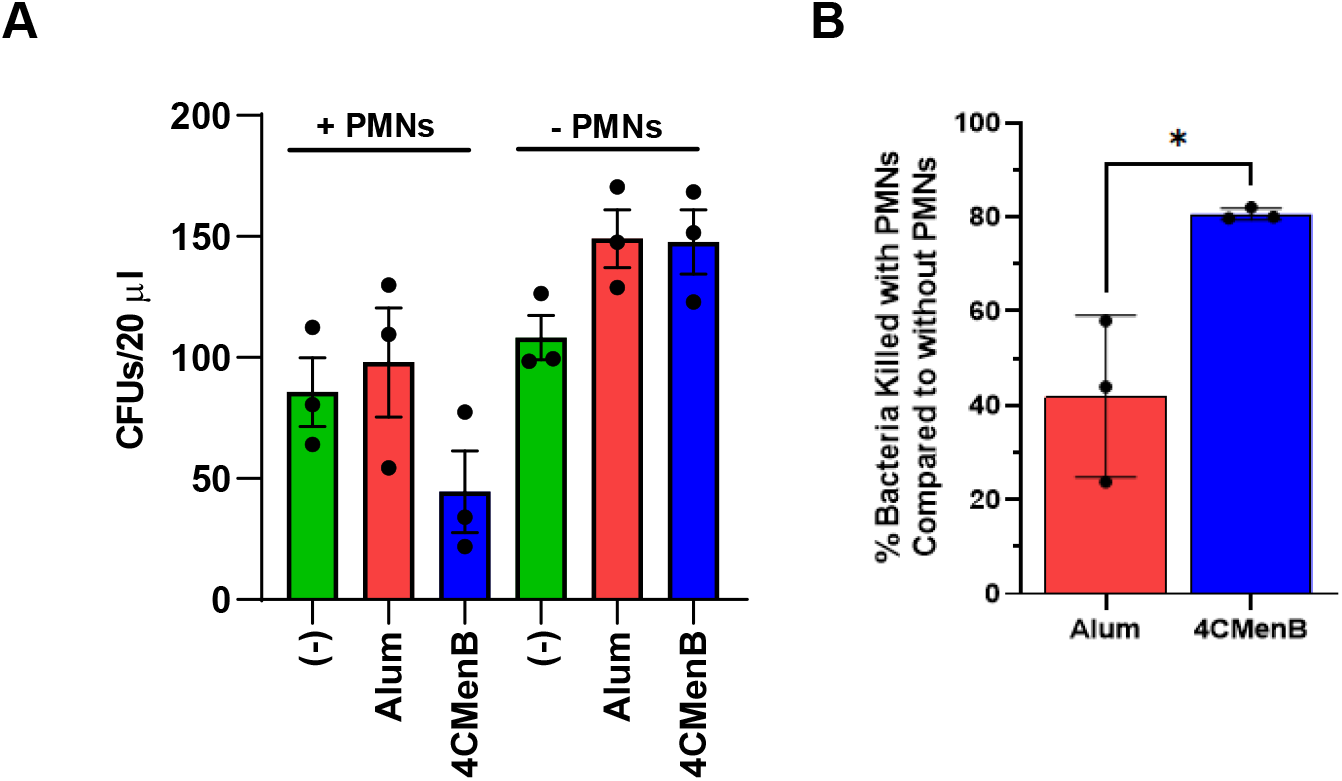
Serum from 4CMenB-vaccinated mice enhances opsonophagocytic killing of *N. gonorrhoeae* by primary human PMNs. A) *N. gonorrhoeae* strain FA1090 was incubated with 2% BSA and 1:10 dilution of pooled serum from 4CMenB-vaccinated (blue) or alum-treated control (red) mice, or no serum (green). 5% C6-depleted human serum that was intact (C’) or heat inactivated (HI-C’) was then added along with primary human PMNs (left side) or buffer control (right side). After 2 hr, CFU were enumerated from the infection mix after overnight growth. (B) The percentage of bacteria killed by OPKA with 4CMenB-vaccinated serum relative to the alum control is presented. Bars indicate the mean ± SD of three independent experiments, with each biological replicate as one data point from a different subject’s PMNs. *, *P* < 0.025, Student’s *t* test.

We conducted the OPKA on serum from 4CMenB-vaccinated and alum-treated mice in the presence or absence of PMNs and with intact C6-depleted NHS in three independent experiments, each with a different subjects’ PMNs. CFUs were enumerated from each condition as in **Fig 7A**. For each mouse serum condition, the CFUs enumerated with PMNs were divided by the CFUs enumerated without PMNs, subtracted from 1 to represent as fraction of bacteria killed in the presence of PMNs, and multiplied by 100%. As shown in **Fig 7B**, there was a statistically significant, reproducible increase in OPKA-mediated killing of strain FA1090 *N. gonorrhoeae* by 4CMenB-vaccinated mouse serum compared with serum from alum-treated mice. However, the magnitude of this effect was approximately two-fold increase in killing, compared with the potency of SBA (see **Fig 6**), which was approximately ten-fold. We conclude that 4CMenB vaccination elicits a measurable but small effect on opsonophagocytic killing of *N. gonorrhoeae* by PMNs.

Taken together, these results show that 4CMenB vaccination of mice elicits antibodies that bind the surface of *N. gonorrhoeae* and promotes SBA and OPKA. These findings align with ongoing investigations into the cross-protective efficacy of OMV-containing vaccines like 4CMenB against *N. gonorrhoeae* and serve as methods for further investigation of these vaccines’ mechanisms of action and correlates of protection.

## Discussion

Preclinical efforts to develop vaccines for gonorrhea warrant the establishment of assays that report the functional, protective activities associated with vaccination, particularly in animal models. Although *N. gonorrhoeae* is a human-restricted pathogen, female mice can be experimentally infected with *N. gonorrhoeae,* and murine models of cervico-vaginal and ascending gonococcal infection are being used to test candidate vaccines against gonorrhea (71). To facilitate the investigation of immune correlates for candidate gonorrhea vaccines in the mouse model, here we report the development of a suite of assays that measure functions associated with vaccine-elicited antibodies in mice: antibody binding to the gonococcal surface, SBA using Ig-depleted pooled normal human serum, and OPKA using C6-depleted pooled normal human serum and primary human PMNs. We report outcomes from these assays using purified antibodies with known gonococcal surface-binding characteristics as proof of concept and as positive controls. We extended these assays to demonstrate that the 4CMenB meningococcal vaccine elicits functional antibodies in mice that bind *N. gonorrhoeae* and elicit cidal activities. These assays can be applied in the context of gonorrhea vaccine development to measure and compare functional outcomes elicited by different vaccine formulations, and review these results in light of clearance rates and colonization loads in individual mice that are immunized and challenged with *N. gonorrhoeae*. Ultimately this information will be used to determine correlates and surrogates of immunity and mechanisms of immune protection in gonorrhea.

This study presents, to the best of our knowledge, the first time a suite of assays has been developed to measure antibody binding, SBA, and OPKA using multiple monoclonal and polyclonal antibodies and normal human serum as the complement source, for evaluating multiple lineages of *N. gonorrhoeae*. As our goal was to incorporate these assays into the workflow for gonorrhea vaccine development, we prioritized rigor and reproducibility considerations. A single stock of each bacterial strain was maintained at −80° C and cultured under the same conditions for each experiment. Antibody binding, SBA, and OPKA assays were conducted in 96-well plates to minimize volumes of bacteria, cells, human serum as complement source, and mouse serum as antibody source. The same positive control antibodies were included in every run of each experiment for comparison between days and human subjects and to ensure all reagents are functioning properly. A single lot of human serum as complement source was purchased in bulk to ensure consistency for the duration of these experiments. The human serum was purchased pre-aliquoted in small volumes to avoid repeated freezing and thawing of complement, which is heat-labile. Human serum was used as complement source for the volume of serum we could obtain, the high complement activity it contains, and to better mimic human infection due to the presence of other serum components that could modulate complement- and antibody-mediated activities, for instance *N. gonorrhoeae* binding of complement-inhibitory proteins like factor H or complement C4b-binding protein (19). For SBA, we used Ig-depleted normal human serum as complement source to remove any potentially cross-reactive endogenous human antibodies from the assay. BSA was added to the SBA and OPKA to minimize background nonspecific or low-affinity antibody binding to Gc and amplify differences in the context of immunization, which may explain differences in the magnitude of our SBA results with those reported by Leduc *et al.* (26). With a number of publications on SBA and OPKA for gonorrhea vaccines, studies that directly compare the SBA and OPKA assays here to ones published by other groups are needed for the field to reach consensus on preclinical vaccine evaluation for *N. gonorrhoeae* (26, 44, 45).

Functional antibody studies for *N. gonorrhoeae* have particular challenges. First, *N. gonorrhoeae* is notoriously phase- and antigenically variable, which can affect day-to-day experimental outcomes. We did not select for any bacterial subpopulations from the variable bacterial stocks, choosing instead for a diverse stock of bacteria in each experiment to reflect the variants that would emerge in human infection (72, 73). These variations could explain why it was not possible to obtain 100% antibody-positive bacteria by imaging flow cytometry. Second, flow cytometry, particularly imaging flow cytometry, is expensive and time-consuming in sample processing, data acquisition, and analysis. ELISA is typically used as a surrogate for antibody binding to bacteria, and is an approach being taken by others in our consortium. However, some antibodies may differ in binding to immobilized epitopes and binding to the bacterial surface, especially if the antigens are membrane proteins that achieve different conformations in the bacterial outer membrane. Future studies will compare the results obtained with imaging flow and ELISA for antigens of interest. Third, while CFU enumeration is an accurate way to monitor antibody-mediated control of bacteria via SBA and OPKA, it takes time and effort to plate and count colonies after overnight growth. The latter could be accelerated using automated colony counting. Our attempts to use the metabolic dye alamarBlue as a surrogate for colony counting for SBA did not yield substantial differences between sensitive and resistant conditions with the low numbers of Gc used for SBA, which agrees with Clow *et al.’s* finding that a luminescence-based alternative for SBA was not as quantitative as CFU enumeration (74). However, these alternative approaches could be optimized in the future to enable higher-throughput analyses of vaccine candidates, as for *N. meningitidis* (75). Fourth, OPKA requires the use of primary human PMNs, but not all investigators can readily source human blood cells. We developed the opsonophagocytic flow cytometry assay with the HL-60 human promyelocyte cell line, which has been used extensively for measuring OPKA against Gram-positive bacteria like *S. pneumoniae,* to avoid the need for primary blood cells. However, results with differentiated HL-60 cells were less consistent than anticipated. Murine PMNs could also be isolated by peritoneal lavage for *in vitro* OPKA to match the murine antibodies elicited by immunization in this preclinical model (76). Fifth, we used C6-depleted NHS as the OPKA complement source to decouple the direct bactericidal effect of serum from phagocyte-mediated killing, but recognize that these activities may be synergistic in the context of infection. Finally, these assays measure the serum antibody response to *N. gonorrhoeae*, which may not accurately reflect the nature of the immune response to the bacteria at the mucosal surfaces they inhabit. To address this issue, we are exploring if the volume and antibody titer in vaginal washes from immunized mice are sufficient for use in the functional antibody studies.

OMV-based meningococcal serogroup B vaccines have been shown to cross-protect humans from gonorrhea infection and hospitalization (20, 21, 23). Our results show that immunization of mice with the 4CMenB meningococcal vaccine elicits antibodies that bind the surface of diverse strains of *N. gonorrhoeae* and promote SBA and OPKA. These findings align with reports that mice vaccinated with 4CMenB are significantly less likely to remain vaginally colonized with *N. gonorrhoeae* (26) and that 4CMenB vaccination elicits antibodies that cross-react with *N. gonorrhoeae* antigens (27). We find it intriguing that the magnitude of the SBA titer did not correlate with the degree of antibody binding to the bacterial surface. Strain MS11, which had the highest SBA titer for 4CMenB-vaccinated serum, also had the lowest bacterial binding, and H041 did not show any specific increase in binding of antibody from 4CMenB-vaccinated serum, yet vaccination elicited SBA 16-fold above the alum-treated control (see **Fig 5** and **Table 2).** Factors that may contribute to this discrepancy include the efficacy of the subclass of antibodies that bind to the bacterial surface to elicit SBA, the presence of other immunoglobulin subtypes in the serum that differentially elicit SBA on different strains (e.g. IgM), the abundance and conformation of the antigen on the surface of each strain, and the intrinsic resistance of different strains to complement-mediated lysis.

Gonorrhea vaccine research has greatly accelerated in recent years, with several candidate vaccines now in the discovery stage (17, 77, 78). Immune parameters including antibody levels elicited by these different vaccine candidates are being measured, and validated functional assays are needed to assess the importance of antibody-mediated protection in experimental challenge models. At this time we do not know which candidate vaccine antigens elicit protective effects, nor do we know what immune parameters are protective (17). Excitingly, many different vaccine candidates are entering preclinical evaluation in the mouse model, including purified proteins, proteins refolded into nanodiscs, peptides presented on virus-like particles, or OMV-based strategies. As results from these candidates become available, we and others will be able to rigorously test the hypothesis that mice that show enhanced clearance of *N. gonorrhoeae* have a stronger functional antibody response in bacterial antibody binding, SBA, and/or OPKA. When these antigens are identified, their contribution to the vaccine-elicited functional antibody response can be confirmed using *N. gonorrhoeae* that is engineered to lack the antigen of interest. Future efforts will be directed at developing functional antibody assays for evaluating serum from vaccinated vs. unvaccinated humans that is collected from clinical trials, such as those ongoing for 4CMenB (including NCT05294588 and (79)).

There are several important outcomes from applying functional antibody assays to gonococcal vaccine discovery. First, we will directly test the hypothesis that vaccine-elicited antibodies are protective against *N. gonorrhoeae* in experimental murine infection. This hypothesis is generally accepted for *N. meningitidis,* where capsular polysaccharides in some serogroups elicit a protective antibody response and where SBA was used to identify the antigens to include in 4CMenB, but remains to be demonstrated for *N. gonorrhoeae.* Second, we will identify the antigens in *N. gonorrhoeae* that elicit serum antibody responses and the protective nature of those responses. These findings can help prioritize antigens to include or exclude in new gonococcal vaccines. For instance, antibodies against the Rmp antigen are detrimental to protective antibody responses against pathogenic *Neisseria* (13, 80). Third, we will test how outcomes in bacterial antibody binding, SBA, and OPKA correlate with one another and with other measures of vaccine efficacy, such as IgG titer and T cell responses. Finally, we will identify parameters that are predictive of a protective response against experimental infection of mice with *N. gonorrhoeae.* These parameters can be used to prioritize gonococcal vaccine antigens for further preclinical testing and optimization, and ultimately for introduction into humans in phased clinical trials. These studies can help achieve the pressing need for vaccines against gonorrhea to mitigate antibiotic resistance and improve global public health.

## Acknowledgements

We thank Asya Smirnov and Stephanie Ragland for their initial contributions in the development of these protocols. We thank Michael Solga of the UVA Flow Cytometry Core Facility (RRID: SCR_017829) for expert advice on imaging flow cytometry, Sanja Arandjelovic for advice on growing and differentiating HL-60 cells, and Sanjay Ram and Peter Rice for the gift of 2C7 monoclonal antibody. We thank all members of the Gonorrhea Vaccine Cooperative Research Center, in particular Margaret Bash, Alex Duncan, and Kathryn Matthias, and Sanjay Ram and Lisa Lewis, for thoughtful discussions on assay development and reporting. This work was supported by NIH U19 AI144180, NIH U01 AI162457, R01 AI097312, and R21 AI157539.

